# Humanized Single Domain Antibodies Neutralize SARS-CoV-2 by Targeting Spike Receptor Binding Domain

**DOI:** 10.1101/2020.04.14.042010

**Authors:** Xiaojing Chi, Xiuying Liu, Conghui Wang, Xinhui Zhang, Lili Ren, Qi Jin, Jianwei Wang, Wei Yang

## Abstract

Severe acute respiratory syndrome coronavirus 2 (SARS-CoV-2) has spread across more than 200 countries and regions, leading to an unprecedented medical burden and live lost. SARS-CoV-2 specific antivirals or prophylactic vaccines are not available. Neutralizing antibodies provide efficient blockade for viral infection and are a promising category of biological therapies. Using SARS-CoV-2 spike RBD as a bait, we have discovered a panel of humanized single domain antibodies (sdAbs). These sdAbs revealed binding kinetics with the equilibrium dissociation constant (KD) of 0.7~33 nM. The monomeric sdAbs showed half maximal inhibitory concentration (IC_50_) of 0.003~0.3 μg/mL in pseudotyped particle neutralization assay, and 0.23~0.50 μg/mL in authentic SARS-CoV-2 neutralization assay. Competitive ligand-binding data suggested that the sdAbs either completely blocked or significantly inhibited the association between SARS-CoV-2 RBD and viral entry receptor ACE2. Finally, we showed that fusion of the human IgG1 Fc to sdAbs improved their neutralization activity by tens of times. These results reveal the novel SARS-CoV-2 RBD targeting sdAbs and pave a road for antibody drug development.

## Introduction

Coronavirus disease 2019 (COVID-19) is caused by infection of emerging severe acute respiratory syndrome-associated coronavirus 2 (SARS-CoV-2) and had been declared by World Health Organization as the first coronavirus pandemic in human history^1^. The severity of COVID-19 symptoms can range from asymptomatic or mild to severe with an estimated mortality rate from less than 2% to up to 10% of patients depending on various factors^2^. SARS-CoV-2 is spreading rapidly and sustainably around the world, urging prompt global actions to develop vaccines and antiviral therapeutics.

SARS-CoV-2 polyprotein shares ~86.15% identity with SARS-CoV (Genbank ID: AAS00002.1) and is classified into the genus betacoronavirus in the family *Coronaviridae*^3^. SARS-CoV-2 is an enveloped, positive-sense, single-stranded RNA virus with a large genome of approximately 30,000 nucleotides in length. The virus-encoded membrane (M), spike (S) and envelope (E) proteins constitute the majority of the protein that is incorporated into SARS-CoV-2 envelope lipid bilayer. The S protein can form homotrimers and protrudes from envelope to show the coronal appearance, invading susceptible cells by binding potential SARS-CoV-2 entry receptor angiotensin converting enzyme 2 (ACE2)^3^. Recently, researchers have figured out the molecular structure of SARS-CoV-2 S protein^4^. It is composed of 1273 amino acids and structurally belongs to the type I membrane fusion protein with two areas S1 and S2. The S1 region mainly includes the receptor binding domain (RBD), while the S2 region is necessary for membrane fusion. The RBD structure determines its binding efficiency with ACE2 and provides an important target for neutralizing antibody recognition.

Single domain antibodies (sdAbs), namely nanobodies, were initially identified from camelids or cartilaginous fish heavy-chain only antibodies devoid of light chains, where antigen-binding is mediated exclusively by a single variable domain (VHH)^5^. Therefore, sdAbs are the smallest fragments that retain the full antigen-binding capacity of the antibody with advantageous properties as drugs, imaging probes and diagnostic reagents^6^. The advantages of short development time, flexible formatting and robust production efficiency make sdAb a powerful means to defeat infectious disease pandemics. For therapeutic purpose, relatively sophisticated humanization techniques have been adopted to modify the camelid-specific amino acid sequences in the framework to their human heavy chain variable domain equivalent, without altering sdAb’s biological and physical properties and reducing species heterogeneity^7^. As SARS-CoV-2 is an emerging human virus, the whole population is susceptible due to the lack of protective antibodies. The existing neutralizing antibodies in convalescent plasma have been adopted as powerful therapeutic alternatives for COVID-19 patients in China. Using a synthetic humanized sdAbs discovery platform, we were able to obtain several high-affinity SARS-CoV-2 RBD targeting sdAbs with desired neutralization activities.

## Results

SARS-CoV-2 makes use its envelope S glycoprotein to gain entry into host cells through binding ACE2. Recent cryo-EM research revealed that the S protein shows an asymmetrical homotrimer with a single RBD in the “up” confirmation and the other two “down”^4^. Antibodies may take advantage of this RBD structure to block virus entry. To enrich for SARS-CoV-2 RBD binding sdAbs, we performed four rounds of biopanning using a lab owned, full synthetic, humanized phage display library with recombinant RBD protein. After phage ELISA identification of 480 clones, a number of sdAbs exhibited an excellent affinity for SARS-CoV-2 RBD. Five distinctive sdAd sequences (1E2, 2F2, 3F11, 4D8 and 5F8) were cloned into a prokaryotic expression vector and recombinant sdAb proteins were purified by nickel-charged sepharose affinity chromatography (Fig. 1a). Humanized sdAbs obtained in this study are about 125 amino acids with a single VHH domain in average molecular weight less than 15KDa (Fig. 1a). The sdAbs consist of three complementarity determining regions (CDRs) as well as four framework regions (FRs). The amino acids in the frameworks have been maximally humanized, except for residues Phe-42 and Gly-52 in framework-2 to maintain proper antigen affinity and best stability^7^.

**Figure 1.**
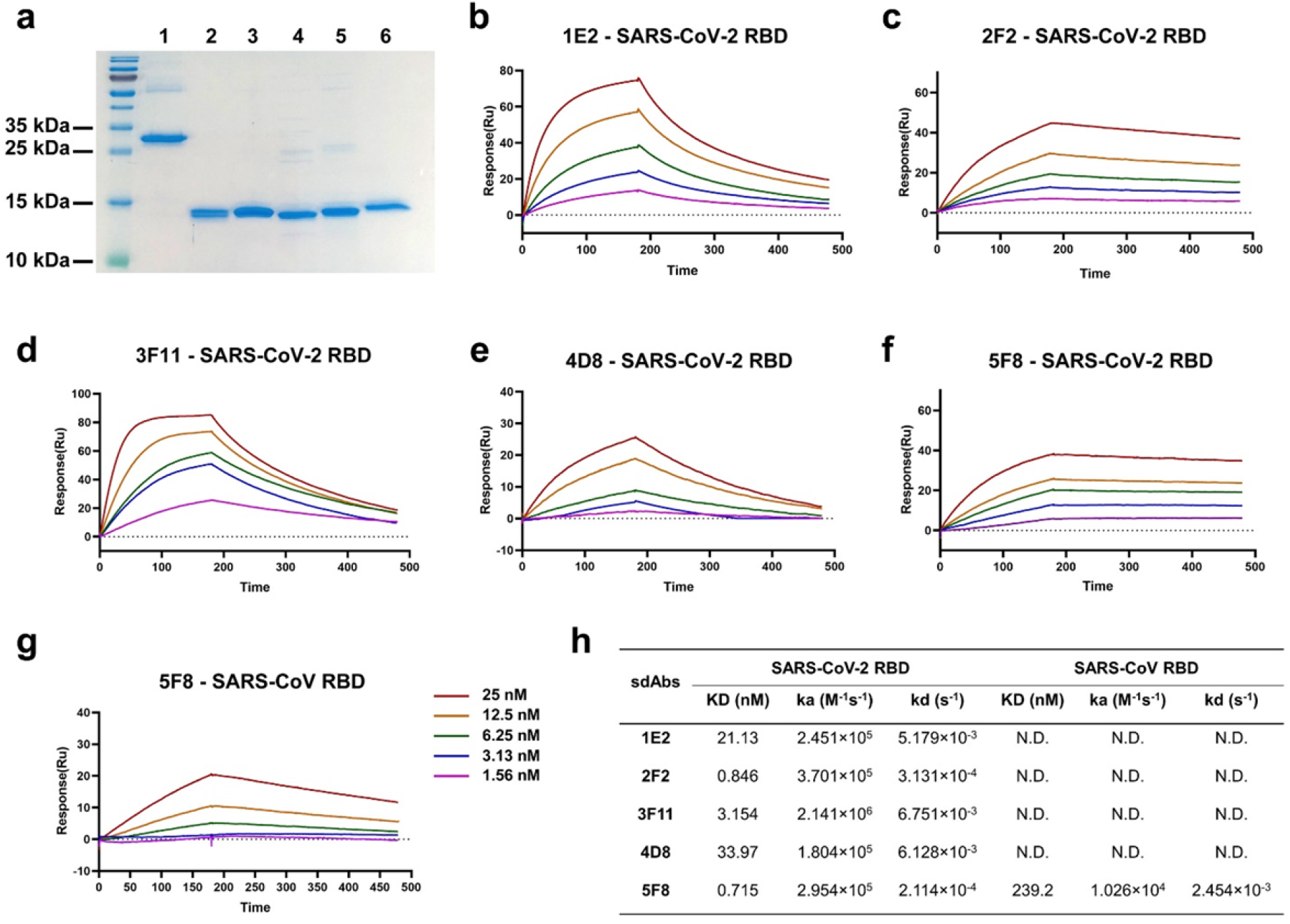
Identification of SARS-CoV-2 RBD binding sdAbs. (**a**) The purified recombinant proteins of SARS-CoV-2 RBD and sdAbs were separated by SDS-PAGE and stained with Coomassie Blue. Lanes: 1, SARS-CoV-2 RBD; 2, 1E2; 3, 2F2; 4, 3F11; 5, 4D8; 6, 5F8. (**b-f**) Five sdAbs binding to SARS-CoV-2 S measured by SPR. Two-fold serial dilutions from 25 nM sdAb injected onto the captured S homotrimer. Kinetic data from one representative experiment were fit to a 1:1 binding model. The profiles are shown for 1E2 (b), 2F2 (c), 3F11 (d), 4D8 (e) and 5F8 (f). (**g**) Kinetics of binding between SARS-CoV S and 5F8. (**h**) Summary of SPR kinetic and affinity measurements. N.D. means not detected.

Surface plasmon resonance (SPR) technology is widely accepted as a golden standard for characterizing antibody-antigen interactions. To determine the kinetic rate and affinity constants, detailed analysis of S antigen-binding to purified sdAb proteins was carried out by SPR. The SARS-CoV-2 or SARC-CoV S protein was immobilized on the surface of Biacore Chip CM5, respectively. Then, various concentrations of purified sdAbs were prepared and injected to pass over the surface. The sensorgram data were fitted to a 1:1 steady-state binding model. SPR results demonstrated that the equilibrium dissociation constant (KD) for the SARS-CoV-2 S protein against sdAbs 1E2, 2F2, 3F11, 4D8 and 5F8 were 21.1 nM, 0.846 nM, 3.154 nM, 33.97 nM and 0.676 nM respectively (Fig. 1b-1f & 1h). However, the sdAbs showed no binding with SARS-CoV S protein, except for the clone 5F8 demonstrating a relatively low affinity with KD n= 239.2 nM (Fig. 1g & 1h). Overall, as monovalent antibody fragment, the sdAbs identified in this study reveals a satisfactory binding performance in a SARS-CoV-2 specific manner.

To further evaluate the neutralization activity of these sdAbs, SARS-CoV-2 S-pseudotyped particle (SARS-CoV-2pp) infectivity assay was first established. Pseudotyped particles are chimeric virions that consist of a surrogate viral core with a heterologous viral envelope protein at their surface, which can be operated in Biosafety Level 2 (BSL-2) and frequently used tool for studying virus entry mechanism and neutralizing antibodies^8^. We observed that all five sdAbs showed inhibition potency of SARS-CoV-2pp infection with IC_50_ (half maximal inhibitory concentration) ranging from 0.003 to 0.3 μg/mL (Fig. 2a). We next tested the neutralization activity of the sdAbs with live SARS-CoV-2 virus (Fig. 2b). Similarly, these sdAbs showed comparable neutralization efficiency, with IC_50_ at approximately 0.2-0.6 μg/mL. Totally, these monovalent sdAbs demonstrated encouraging neutralization activity against both pseudotyped and authentic virus, although the neutralization potency is not completely matched (Fig. 2c). This phenomenon was normally reported in Middle East Respiratory Syndrome coronavirus (MERS-CoV) neutralizing antibodies and may be likely explained by the difference in sdAb recognized RBD spatial epitope or the steric hindrance formed by antigen-antibody complex^9,10^.

**Figure 2.**
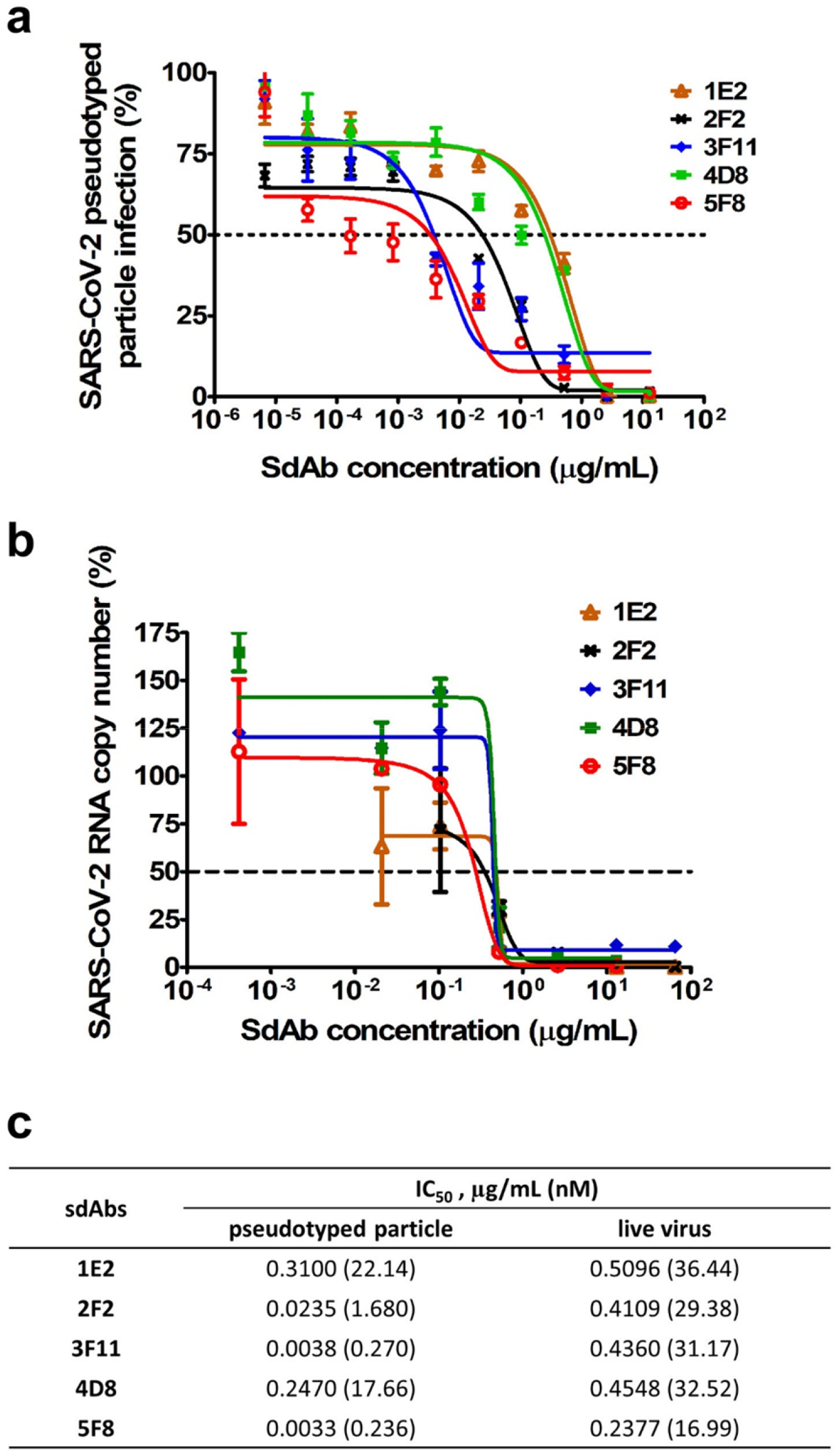
Neutralization of SARS-CoV-2 by RBD-specific sdAbs. (**a**) Neutralization of 5 sdAbs against SARS-CoV-2pp. SARS-CoV-2pp was pre-incubated with 5-fold serially diluted sdAbs before inoculation of Calu-3 cells. At 48 h post infection, luciferase activities were measured, and percent neutralization was calculated. (**b**) Determination of neutralization activities of 5 sdAbs against live SARS-CoV-2. Absolute quantification of SARS-CoV-2 RNA copy number in culture supernatants was performed using real time RT-PCR method, and percent neutralization was calculated. (**c**) Summary of the half maximal inhibitory concentration (IC_50_) values of the 5 sdAbs against both SARS-CoV-2pp and live virus.

Within SARS-CoV-2 RBD, the receptor binding motif (RBM) directly contacts ACE2. Recent report demonstrating that SARS-CoV-2 uses ACE2 as its receptor with a much stronger affinity (10- to 20-fold higher) than SARS-CoV^4^. To determine whether sdAbs targeted different antigenic regions on the SARS-CoV-2 RBD surface, we performed a competition-binding assay using a real-time biosensor (Fig. 3). We tested all five sdAbs in a competition-binding assay in which human ACE2 was attached to a CM5 biosensor. Compared with a non-related isotype control sdAb (Fig. 3a), addition of 1E2, 3F11 and 4D8 completely prevent binding of SARS-CoV-2 RBD to ACE2 (Fig. 3b, 3d, 3e). Whereas, sdAbs 2F2 and 5F8 could partially compete the RBD/receptor association (Fig. 3c & 3f). These data suggested that these sdAbs can be divided into RBM targeting or non-RBM targeting groups though it is not directly associated with virus neutralization activity.

**Figure 3.**
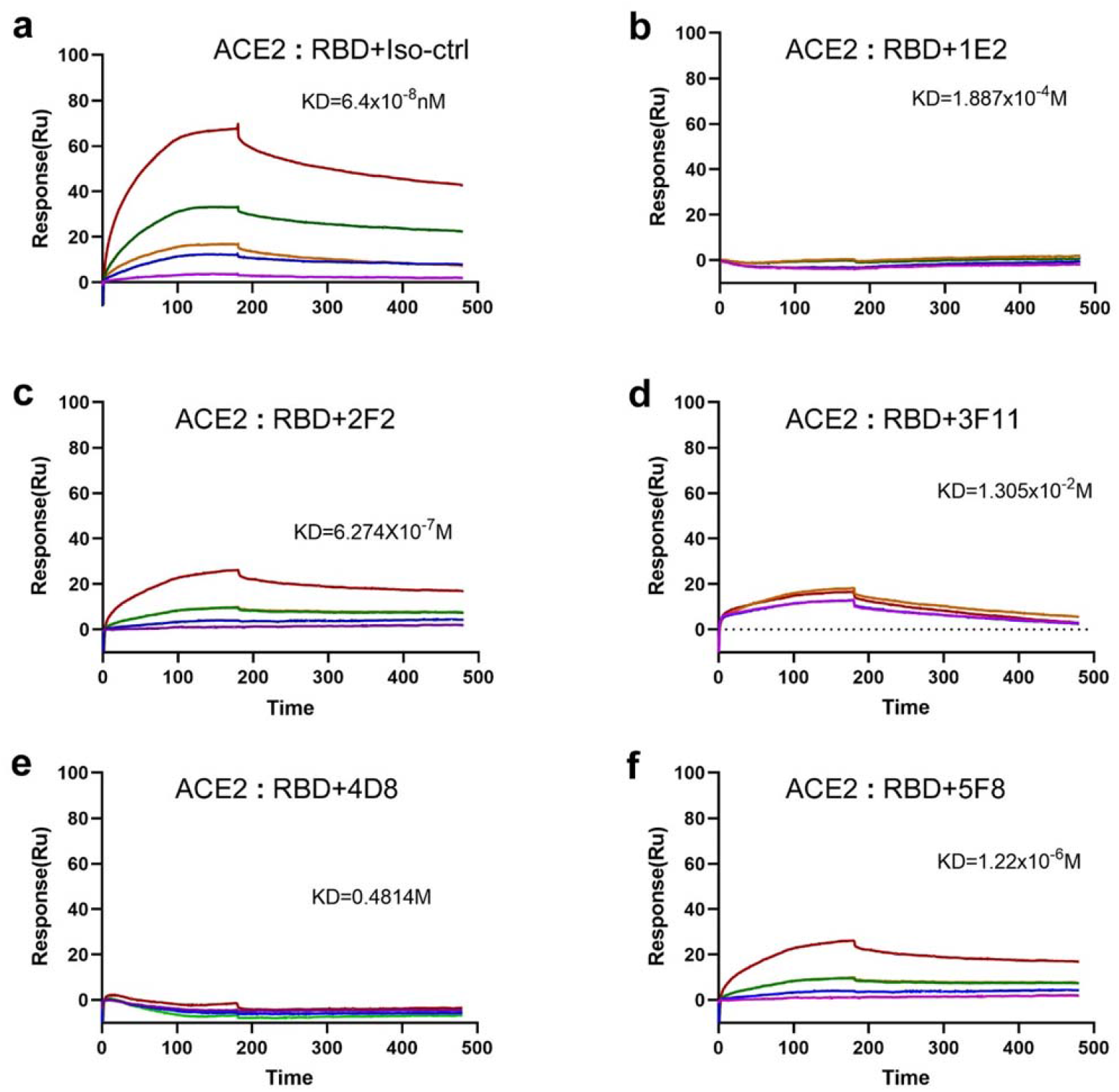
Interference of the ACE2-RBD interaction by the sdAbs. The recombinant human ACE2 protein was immobilized on CM5 chip using a BIAcore T200 machine and tested for the binding with gradient concentrations of SARS-CoV-2 RBD that were diluted in 100nM sdAbs, including an isotype control sdAb (**a**), 1E2 (**b**), 2F2 (**c**), 3F11 (**d**), 4D8 (**e**) and 5F8 (**f**).

SdAbs can be readily fused to human Fc-domain to overcome the limitations of the monovalent sdAbs, such as the short blood residential time and lacking antibody-dependent cell-mediated cytotoxicity and complement dependent cytotoxicity^11^. In addition, bivalent sdAbs can be obtained via the disulfide bond formation in Fc hinge area, which was reported to significantly increase sdAb’s activity^12^. To further explore the possibility of sdAb-based antiviral therapeutics and enhance neutralization activity, we constructed human heavy chain antibodies by fusing the human IgG1 Fc region to the C-terminus of sdAbs (Fig. 4a & 4b). These Fc fusion sdAbs were produced in mammalian cells with suspension yields around 25-50 μg per milliliter in shaking flask. The supernatant samples were analyzed in both reducing and non-reducing conditions in Western blot using an anti-human IgG to detect Fc. As shown in Fig. 4c, the size of the constructed intact sdAb-Fc is around 80 KDa in the non-reducing condition, but a 40 KDa monomer was observed by prior treatment in reducing condition to break disulfide bonds. This suggests a correct expression and secretion of heavy chain antibodies in consistence with our design. Neutralization assay results showed that genetic fusion of human Fc significantly increased the neutralization activity of these sdAbs for 10- to 80-fold in molar concentration of IC_50_ using the SARS-CoV-2pp entry assay (Fig. 4d). Importantly, several Fc-fused sdAbs demonstrated potency with IC_80_ at sub-nanomolar level (Fig. 4d). Finally, we showed that some of the sdAbs are suitable for immunofluorescence staining (Supplementary Fig. 1) and Western blot to detect ectopically expressed SARS-CoV-2 S protein (Supplementary Fig. 2).

**Figure 4.**
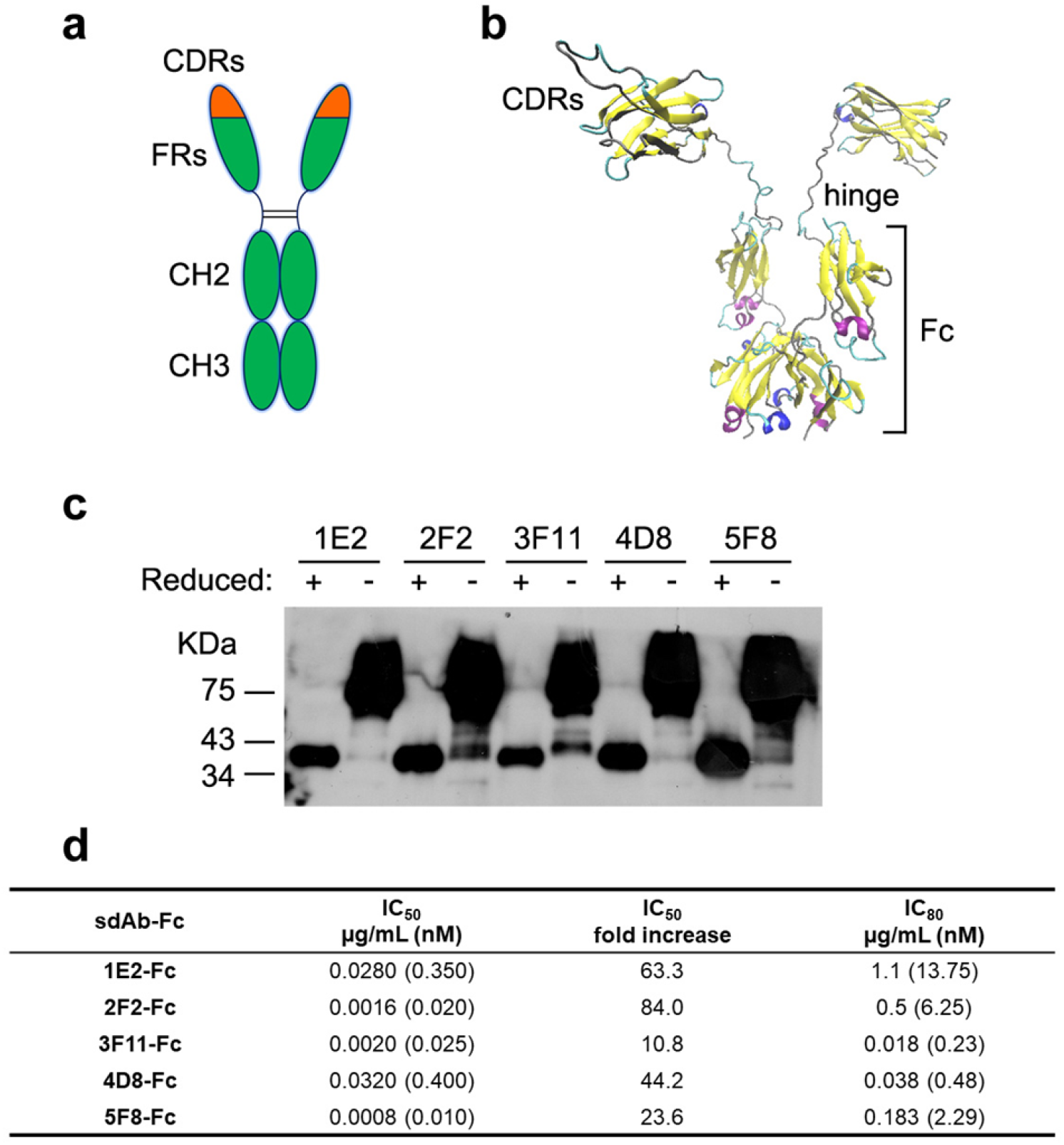
Inhibition of SARS-CoV-2 entry by Fc-fused sdAbs. (**a**) Representation of the human IgG1 Fc-fused sdAbs in this study. SdAb-Fc fusion construction generates a bivalent molecule with an approximate molecular weight of 80 kDa. (**b**) Homology modeling of the bivalent 5F8-Fc molecule with SWISS-MODEL server (https://swissmodel.expasy.org)^22^. The template structure for 5F8 modeling was based on a humanized camelid sdAb in the PDB database (3EAK). The structure is depicted as cartoons and colored with secondary structure. Three CDRs, hinge region and Fc were indicated. (**c**) Five Fc-fused sdAbs were analyzed by Western blot with gradient SDS-PAGE in reducing (with β-mercaptoethanol) or non-reducing (without β-mercaptoethanol) condition. (**d**) Summary of IC_50_ and IC_80_ values of Fc-fused sdAbs neutralization against SARS-CoV-2pp. IC_50_ fold increases versus the corresponding monovalent sdAbs were calculated.

## Discussion

Given the disease severity and rapid global spread of COVID-19, there is an urgent need for development of vaccines, monoclonal antibodies, and small-molecule direct-acting antiviral medications. Neutralizing antibodies directly target viral envelope protein, precisely block the virus-receptor association, and inhibit virus entry through a variety of molecular mechanisms. In this study, we isolated and characterized several humanized neutralizing sdAbs that exhibit one- to two-digit nanomolar or even sub-nanomolar IC_50_ against SARS-CoV-2 using both pseudotyped and infectious viruses. SdAbs have been investigated as important therapeutic alternatives against viral infection because of their high yield, low cost and intrinsic stability. For MERS-CoV, neutralizing sdAbs were isolated from immunized dromedary camels or llamas and demonstrated IC_50_ value between 0.001-0.003μg/mL with low KD values (0.1-1 nM)^13,14^. Comparable inhibition efficiency on SARS-CoV-2pp and affinity kinetics were obtained for the sdAbs 2F2, 3F11 and 5F8 identified in this study using a non-immune library, which can speed up the discovery of neutralizing antibodies in an emergent outbreak. With further optimization and increase of library size and diversity, the synthetic sdAb library technology will promote the discovery speed of powerful therapeutic antibodies^15,16^.

FDA approved the first sdAb-based medicine for adults with acquired thrombotic thrombocytopenic purpura in 2019^17–20^. Considering the cost and potential risks of full human antibody in some viral diseases, such as dengue virus infection, sdAb fragments are a novel category of therapeutic molecules and can be readily reconstructed in a tandemly linked way to increase their blood residential time, biological activity, and eliminate underlying concerns about antibody-dependent enhancement (ADE) of viral infection^21^. In addition to being used as an injectable drug, the stable sdAbs can be also developed into aerosolized inhalations and disinfection products for the prevention of COVID-19. Besides, prior to the success of COVID-19 vaccines, the construction of sdAb-based adenovirus or adeno-associated virus gene therapy might provide long-term passive immune protection in vulnerable population, health care workers, or in severely affected areas. Since the mature COVID-19 animal models have not been developed, this study did not involve *in vivo* studies. As a next step, the crystal structure analysis of antigen-antibody complexes will be put on the agenda. In conclusion, the discovered neutralizing antibodies in this study could lead to new specific antiviral treatments and shed light on the design and optimization of COVID-19 vaccines.

## Methods

### Cells and reagents

The Vero (African green monkey kidney), HEK293T (human kidney epithelial), 293F, Calu-3 (human lung adenocarcinoma) cells were obtained from China Infrastructure of Cell Line Resource (Beijing, China) and maintained in Dulbecco’s modified Eagle’s medium (DMEM, ThermoFisher, Waltham, MA, USA) supplemented with 2-10% fetal bovine serum (FBS, ThermoFisher), non-essential amino acid, penicillin and streptomycin. Recombinant proteins were purchased from Sino Biological (Beijing, China) for SARS-CoV-2 RBD (40592-V05H, 40592-V08B), SARS-CoV-2 spike (40589-V08B1), SARS-CoV spike (40150-V08B1), ACE2 (10108-H08H). Antibodies were obtained from ThermoFisher for anti-His-HRP, anti-human IgG-HRP, anti-His-488 and anti-CM13.

### Library design and construction

A synthetic sdAb phage display library was used for the screening of SARS-CoV-2 neutralizing antibodies. To minimize a possible antigenic effect from camelid sequences, sdAb frameworks for library construction were determined according to a universal humanized scaffold architecture^7^. Briefly, residues in frames 1, 3 and 4 were mutated based on human heavy chain VH in maximum. In frame 2, humanization of residues at positions 49 and 50 was adopted to increase stability of sdAbs, whereas residues 42 and 52 are maintained in camelid due to their critical impact on antigen affinity and/or stability. For the design of variable regions, we analyzed a robust CDR repertoire from immune or naïve llama VHH clones. A synthetic diversity was introduced in the three CDRs by random nucleotide incorporation with cysteine and stop codon avoided. A constant length of 8 amino acids was selected for CDR1 and CDR2, and 18 amino acids for CDR3. Frameworks and CDRs were assembled by overlapping polymerase chain reaction (PCR) ensuring each unique CDR recombined in the assembled molecules. Diversified sdAb mixture was cloned in phagemid vector fADL-1 (Antibody Design Labs, San Diego, CA, USA) using SfiI/BglI sites with the PelB peptide leader sequence fused with the sdAbs at N-terminus. Massive electroporation was carried out using E. coli TG1 cells. More than a thousand agar petri dishes (140 mm) were plated to ensure enough size of the library. Quality control was carried out by sequencing more than 1000 clones, and the error rate and diversity were calculated.

### Antibody selection by phage display

Screening for SARS-CoV-2 RBD targeting antibodies was performed by panning in both immunotubes and native condition using a proprietary full-synthetic library of humanized sdAbs with high-diversity, according to a standard protocol. Briefly, for the 2^nd^ and 4^th^ panning rounds, the purified SARS-CoV-2 RBD protein was coated on Nunc MaxiSorp immuno tubes (ThermoFisher) at around 5μg/mL in PBS overnight. For the 1^st^ and 3^rd^ panning rounds, RBD protein was first biotinylated with EZ-Link™ Sulfo-NHS-LC-Biotin (ThermoFisher) and then selected with streptavidin-coated magnetic Dynabeads™ M-280 (ThermoFisher). The tubes or beads were blocked using 2% w/v skimmed milk powder in PBS (MPBS). After rinsing with PBS, about 1×10^13^ phage particles were added to the antigen-coated immuno tube or biotinylated antigen in the presence of 2% MPBS, incubated for 2 h shaking (30 rpm) at RT. Unbound phages were washed with PBS Tween 0.1% (10 times) and PBS (10 times), while bound phage were eluted with 0.1M Glycine-HCl (pH=3.0). Eluted phages were neutralized by adding 1M Tris-Cl pH 9.0 and used for infection of exponentially growing E. coli TG1. After 4 rounds of panning, phage ELISA identification was performed with 480 individual colonies using Anti-CM13 antibody [B62-FE2] (HRP). The absorbance was measured using a SpectraMax M5 plate reader from Molecular Divices (San Jose, CA, USA). The positive clones were sent for sequencing, and representative sdAb sequences were chosen for protein expression.

### Expression and purification of sdAbs

Full-length sequences of selected sdAbs were PCR amplified and cloned into the NcoI/XhoI sites of the pET28b (Novagen, Sacramento, CA, USA) and transformed into BL21(DE3) chemically competent E. coli. A single colony was picked to inoculate 10 ml of LB media containing Kanamycin (100 μg/mL) and incubated at 37°C on an orbital shaker overnight. This preculture was diluted 1:100 in 400 mL of LB media containing Kanamycin (100 μg/mL) and grown at 37°C until the OD_600_ nm reached 0.4. The expression of recombinant sdAbs was induced by adding IPTG to a final concentration of 0.3 mM after culture has reached OD_600_=0.5-0.6 and grown over night at 20°C. The sdAbs with a His-tag fused at C-terminus were purified over Ni Sepharose 6 Fast Flow (GE Healthcare, Boston, MA, USA) and eluted with 400 mM imidazole. Affinity purified sdAbs were dialyzed against PBS to eliminate imidazole.

### Affinity measurement and competition-binding Study

The surface plasmon resonance experiments were performed at room temperature using a BiaCore T200 with CM5 sensor chips (GE Healthcare). The surfaces of the sample and reference flow cells were activated with a 1:1 mixture of 0.1 M NHS (N-hydroxysuccinimide) and 0.1 M EDC (3-(N,N-dimethylamino) propyl-N-ethylcarbodiimide) at a flow rate of 10 μL/min. The reference flow cell was left blank. All the surfaces were blocked with 1 M ethanolamine, pH 8.0. The running buffer was HBS-EP (0.01M HEPES, pH 7.4, 150 mM NaCl, 3 mM EDTA, 0.05% surfactant P20).

For binding affinity assays, the SARS-CoV-2 or SARS-CoV S protein was diluted in 10mM sodium acetate buffer, pH5.5, and was immobilized on the chip at about 300 response units. Antibodies 1E2, 2F2, 3F11, 4D8 and 5F8 at gradient concentrations (0, 1.56nM, 3.125nM, 6.25nM, 12.5nM, 25nM) were flowed over the chip surface. After each cycle, the sensor surface was regenerated with10mM glycine-HCl pH 2.5. The data were fitted to a 1:1 interaction steady-state binding model using the BIAevaluation 1.0 software.

For competition-binding assays, the ACE2 protein was diluted in 10mM sodium acetate buffer, pH4.5, and was immobilized on the chip at about 650 response units. For the analyses, RBD protein was diluted in HBS-EP buffer or HBS-EP buffer with 100nM antibody (1E2, 2F2, 3F11, 4D8 or 5F8). The RBD in different buffer at gradient concentrations (0, 6.25 25, 100, 400nM) was flowed over the chip surface. After each cycle, the sensor surface was regenerated with 10mM glycine-HCl pH 2.5. The binding kinetics was analyzed with the software of BIAevaluation using a 1:1 binding model.

### Production of SARS-CoV-2 spike pseudotyped particle (SARS-CoV-2pp) and virus entry assay

To produce SARS-CoV-2pp, HEK293T cells were seeded 1 day prior to transfection at 2.5×10^6^ cells in a 10-cm plate. The next day, cells were transfected using Lipofectamine 2000 (ThermoFisher). The plasmid DNA transfection mixture (1 ml) was composed of 15 μg of pNL-4.3-Luc-E^-^R^-^ and 15 μg of pcDNA-SARS-CoV-2-S that was purchased from Sino Biologicals and reconstructed by deletion of 18 amino acid cytoplasmic tail. A nonenveloped lentivirus particle (Bald virus) was also generated as negative control. Sixteen hours after transfection, the media was replaced with fresh media supplemented with 2% FBS. Supernatants containing SARS-CoV-2pp were typically harvested 36–48 h after transfection and then filtered through a syringe filter. The typical relative luminometer units for SARS-CoV-2pp were between 10^6^ and 10^7^. To conduct the virus entry assay, Vero E6 or other cells were seeded in a 96-well plate at 1 day prior to transduction. The next day, 100 μL of supernatant containing SARS-CoV-2pp was added into each well in the absence or presence of serially diluted sdAbs or human IgG1 Fc-fused sdAbs. Forty-eight hours after transduction, cells were lysed in 100 μL of passive lysis buffer and 50 μL lysate was incubated with 100 μL of luciferase assay substrate according to the manufacturer’s instructions (Promega, Madison, WI, USA).

### SARS-CoV-2 neutralization assay

SARS-CoV-2 was isolated from bronchoalveolar lavage specimens from a COVID-19 patient as described previously^23^. The viral titers were determined by using tissue culture infective dose (TCID50). For antibody neutralization assay, Vero cells were seeded in 96-well plates at 1 day prior to infection. Serially diluted sdAbs were mixed with SARS-CoV-2 at a multiplicity of infection (MOI) of 0.05 and incubated at 37°C for 1h. The antibody-virus mixture was incubated on Vero cells at 37°C for 1h. Unbound SARS-CoV-2 virions were removed by washing cells with fresh medium, then incubated for 24 h at 37 °C. The culture supernatants were collected for viral nucleic acid quantification. Viral nucleocapsid gene RNA quantification was carried out by TaqMan real-time RT-PCR as reported with plotted standard curves using in vitro transcribed RNA.

### Production of human IgG1 Fc fusion sdAbs

The sequences of selected sdAbs were cloned into a mammalian expression vector under the control of hEF1-HTLV promotor and fused with N-terminal interleukin-2 signal peptide and C-terminal Fc region, comprising the CH2 and CH3 domains of human IgG1 heavy chain and the hinge region. Maxipreped plasmids were transiently transfected into 293-F cells (Thermofisher) and the cells were further cultured in suspension for 6 days before harvesting antibody-containing supernatant. Fc-fused sdAbs were prepared with prepacked HiTrap® Protein A HP column (GE Healthcare). The produced Fc-fusion protein was analyzed by SDS-PAGE and the Western blot using standard protocols for dimerization, yield and purity measurement. The primary antibody used for Western blot was a horseradish peroxidase conjugated goat anti-mouse IgG (Sigma-Aldrich, St. Louis, MO, USA).

### Immunofluorescence microscopy and Western blot

Cultured 293T cells on coverslips were transfected with either SARS-CoV-2 S expression plasmid or empty vector for 24 h and then fixed using 4% paraformaldehyde for 15 min at room temperature, permeabilized with 0.1% Triton X-100 (Sigma-Aldrich) in PBS for 10 min. The cells were then incubated with each sdAb overnight at 4°C. After three washes with PBS, the cells were incubated with Alexa Fluor 488–conjugated 6x-His Tag monoclonal antibody (ThermoFisher) for 1 hour at room temperature. The nuclei were stained with DAPI (1:10,000) diluted in PBS for 5 min and mounted with an antifade reagent (ThermoFisher). Images were acquired with a Leica TCS SP5 confocal microscope system.

For Western blot, 293T cells in 6-well plate were transfected with SARS-CoV-2 S, SARS-CoV-2 S or empty vector individually. Twenty-four h post transfection, cell lysates were prepared, and the samples were boiled with 2× SDS loading buffer and loaded onto a 10% polyacrylamide gel. After electrophoresis, the separated proteins were transferred onto a nitrocellulose membrane (Bio-Rad, Hercules, CA, USA). The resulting blots were probed with a sdAb as primary antibody and an HRP-linked 6x-His Tag antibody (Thermofisher, HIS.H8) as the secondary antibody. The ECL reagent (Amersham Biosciences, Piscataway, NJ, USA) was used as the substrate for detection.

### Statistical analysis

Data were analyzed using GraphPad Prism 6.01 (GraphPad Software, San Diego, CA, USA). The values shown in the graphs are presented as means ± SD. IC_50_ was calculated based on curve fitness with nonlinear regression and showed by one representative result from at least two independent experiments.

## Acknowledgments

Funding: This work was supported by CAMS Initiative for Innovative Medicine Grant 2016-I2M-3-020.

## Author contributions

W.Y., X.C., L.R. Q.J. and J.W. designed experiments and interpreted the data. W.Y., X.C., X.L., C.W. and X.Z. performed experiments and analyzed the data. W.Y. conceived the study, supervised the work, and wrote the paper. All authors read and approved the final manuscript.

