## Supplemental Figures for "Humanized Single Domain Antibodies Neutralize SARS-CoV-2 by Targeting Spike Receptor Binding Domain"

**Supplementary Information**

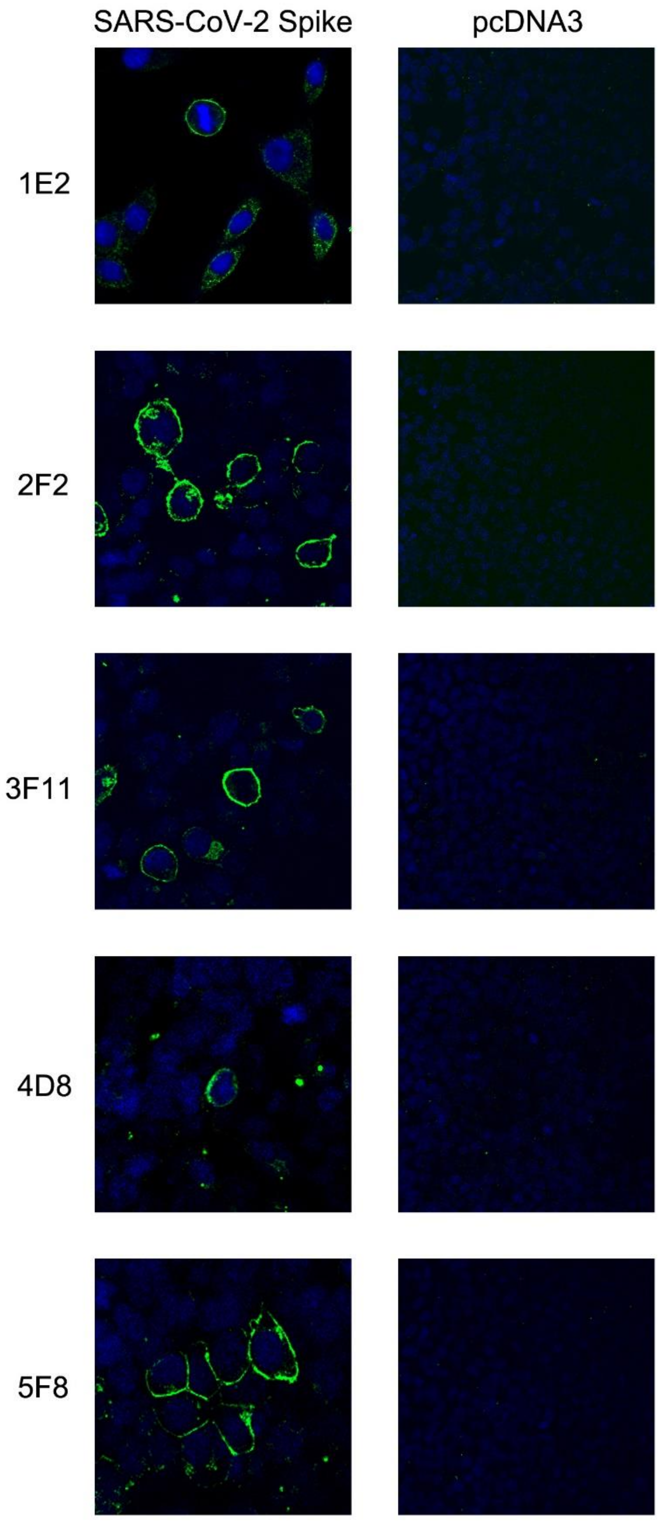


**Supplementary Fig. 1.** Staining of the transfected SARS-CoV-2 S protein in 293T cells with the sdAbs identified in this study. Overexpressed SARS-CoV-2 S protein was clearly localized on 293T cell plasma membrane.

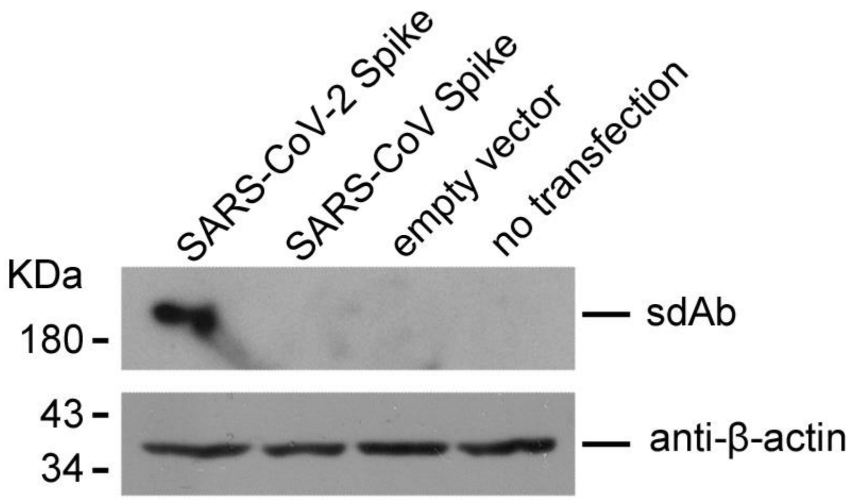


**Supplementary Fig. 2.** Western blot analysis shows specific recognition of SARS-CoV-2 S protein by the sdAb 2H9.
